# Bile acids target proteolipid nano-assemblies of EGFR and phosphatidic acid in the plasma membrane for stimulation of MAPK signaling

**DOI:** 10.1101/335604

**Authors:** Hong Liang, Mary K. Estes, Huiling Zhang, Guangwei Du, Yong Zhou

## Abstract

Bile acids are critical biological detergents in the gastrointestinal tract and also act as messengers to regulate a multitude of intracellular signaling events, including mitogenic signaling, lipid metabolism and endo/exocytosis. In particular, bile acids stimulate many receptors and ion channels on the cell surface, the mechanisms of which are still poorly understood. Membrane-associating proteins depend on the local spatial distribution of lipids in the plasma membrane (PM) for their function. Here, we report that the highly amphipathic secondary bile acid deoxycholic acid (DCA), a major constituent in the human bile, at doses <1μM enhances the nanoclustering of phosphatidic acid (PA) but disrupts the local segregation of phosphoinositol 4,5-bisphosphate (PIP_2_) in the basolateral PM of the human colorectal adenocarcinoma Caco-2 cells. PA has been shown to be a key structural component of the signaling nano-domains of epidermal growth factor receptor (EGFR) on the cell surface. We show that DCA promotes the co-localization between PA and EGFR, the PA-driven EGFR dimerization/oligomerization and EGFR signaling. Depletion of PA abolishes the stimulatory effects of DCA on the EGFR oligomerization and signaling. This effect occurs in the cultured Caco-2 cells and the *ex vivo* human intestinal enteroids. We propose a novel mechanism, where the amphiphilic DCA monomers alter the nano-assemblies of anionic phospholipids and in turn change the dynamic structural integrity of the lipid-driven oligomerization of cell surface receptors and their signal transduction.

## Introduction

Bile acids are synthesized in the liver, stored in the gallbladder and secreted into the small intestine as a component of the enterohepatic circulation [1]. As biological detergents, the bile acids above their critical micelle concentrations (CMCs) form micelles to emulsify fat-soluble molecules to facilitate digestion of fatty acids and lipids in food. The bile acid micelles also act as vehicles to incorporate endogenous and exogenous hydrophobic waste for excretion. At doses well below their CMCs, the bile acids impact cell signaling cascades, including lipid metabolism, mitogenic signaling, ion channel activation, as well as protein trafficking [2]. These signaling effects contribute to the bile acid-induced pathophysiological conditions, including cholestasis, inflammatory bowel disease and cancer in the GI tract [3, 4]. The ability of bile acids to stimulate the nuclear receptors partially contributes to the bile acid-induced changes in cell function and has been studied extensively [5]. However, how the bile acids activate a variety of cell surface receptors is still poorly understood.

Most membrane-associating surface receptors and ion channels dimerize / oligomerize, which is mediated by the heterogeneous distribution of lipids in the plasma membrane (PM), for their function [6-9]. Although the most well-characterized properties of the bile acids have been their amphiphilicity and their ability to associate with lipids as micelles, the typical physiologically effective doses for the bile acids are well below their CMCs. This implies that interactions between the bile acids and the lipids may not contribute to the biological effects of the bile acids. However, the latest *in vitro* studies using isolated natural PM and synthetic vesicles/planar bilayers show that the bile acid monomers at doses well below their CMCs intercalate in the lipid bilayers and alter the highly dynamic and transient nanometer-scale lateral spatial distribution of the lipids [10-12], without changing the global membrane properties, such as overall membrane permeability and lipid solubility. Thus, it is possible that the bile acids stimulate the cell surface receptors via modulating the structural integrity of the lipid-mediated nano-assemblies of the membrane receptor on the cell surface. Here, we used super-resolution electron microscopy (EM) combined with quantitative spatial analysis to systematically screen how the secondary bile acid, deoxycholic acid (DCA), at doses <1μM modulated the nano-segregation of various PM lipids in the basolateral PM of human colorectal adenocarcinoma Caco-2 cells. We then used epidermal growth factor receptor (EGFR) as an example to test the potential consequence of the DCA-induced dimerization/oligomerization and the signaling of EGFR in Caco-2 cells and *ex vivo* human intestinal enteroids.

## Results

### DCA monomers alter the spatiotemporal organization of plasma membrane phospholipids

Bile acid micelles are effective biological detergents capable of solubilizing lipids and elevating membrane permeability. However, it is not clear how the amphiphilic bile acid monomers at physiological levels potentially influence the much more dynamic nanoscale lipid organization in the PM. As local nanoscale aggregation of the lipids directly participates in the dimerization/oligomerization of the cell surface receptors, the lateral mobility of PM lipids is important for cell signaling [7, 9, 13]. We, thus, tested how DCA influenced the spatial distribution of acidic lipids in the Caco-2 basolateral PM.

We ectopically expressed GFP-tagged lipid binding domains in Caco-2 cells: GFP-LactC2 to label phosphatidylserine (PS) [14], GFP-PASS to label phosphatidic acid (PA) [15], GFP-PH-PLCδ to label phosphoinositol 4,5-bisphosphate (PIP_2_) [16], GFP-PH-Akt to label phosphoinositol 3,4,5-trisphosphate (PIP_3_) [17] or GFP-D4H to label cholesterol [18]. To eliminate the effect of high levels of bile acids typically present in bovine serum [19], we pre-serum-starved Caco-2 cells for 2h using serum-free EMEM medium before DCA treatment. The spatial organization of each lipid probe was evaluated on intact basolateral PM sheets prepared from the Caco2 cells using EM-immunogold labeling (Fig. S1A and B) and unvariate K-function spatial analysis expressed as *L(r)-r* [20, 21]. Fig. S1C shows a plot of *L*(*r*)-*r* vs. *r*, where *L*(*r*)-*r* is the extent of nanoclustering and *r* is the length scale in nanometers (nm). *L*(*r*)-*r* values above the 99% confidence interval (99% C.I.) indicate statistically significant clustering, with larger *L*(*r*)-*r* values corresponding to more extensive clustering. The peak *L*(*r*)-*r* value, *L*_*max*_, of each curve summarizes the clustering data (Fig. 1A).

**Figure 1.**
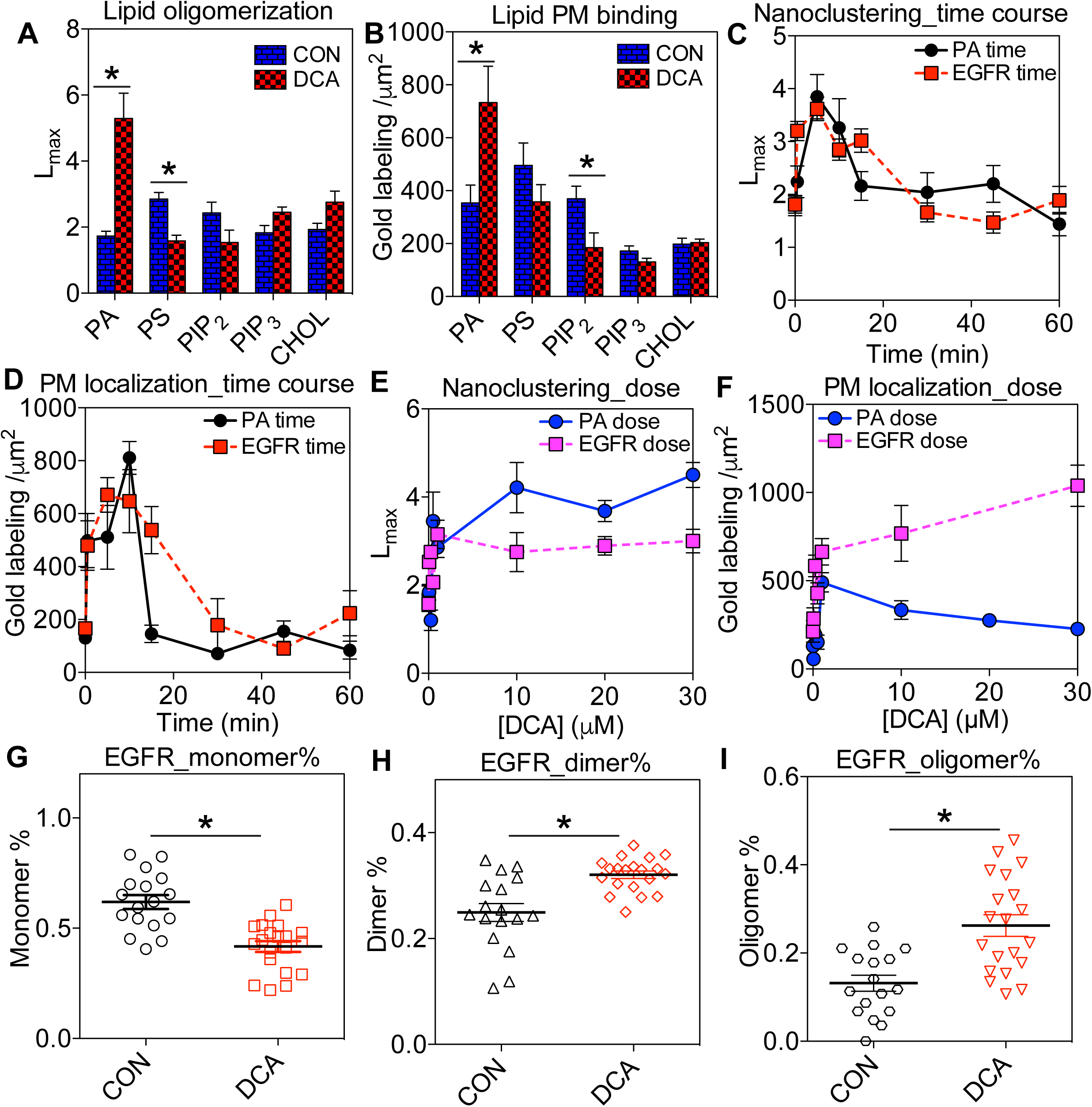
DCA alters the dynamic spatial distribution of various lipids and EGFR in Caoc-2 basal PM. (**A**) Caco-2 cells ectopically expressing a GFP-tagged lipid-binding domain were treated with 1μM DCA for 5 minutes. Intact basolateral PM sheets of Caco-2 cells were then attached to EM grids and immunolabeled with 4.5nm gold nanoparticles conjugated to anti-GFP antibody. Gold particles were imaged via transmission EM at 100,000X magnification. Spatial distribution of gold particles within a 1μm^2^ PM area was quantified using univariate K-function. *L*_*max*_ values above 1 indicate statistically meaningful clustering, whereas *L*_*max*_ values below 1 indicate spatial uniformity. (**B)** Number of gold particles within the same 1μm^2^ PM area was counted as an estimate of lipid level in the PM. (**C** and **D**) Caco-2 cells expressing either GFP-PASS (to specifically tag PA) or EGFR-GFP were exposed to 1μM DCA for various time points (0-60 minutes) before immunogold labeling. Extent of univariate spatial distribution (C) and number of gold labeling (D) of GFP-PASS or EGFR-GFP were calculated as described above. (**E** and **F**) Caco-2 cells ectopically expressing GFP-PASS or EGFR-GFP were exposed to increasing concentrations of DCA (0-30μM) for 5 minutes before immunogold labeling. Spatial distribution (E) and number of gold particles within a 1μm^2^ PM area (F) were calculated. Changes in EGFR monomer (**G**), dimer (**H**) and oligomer (**I**) populations without / with 1μM DCA were calculated using the spatial analysis data. At least 15 PM sheets from individual cells were analyzed in each condition. Statistical significance of the univariate clustering analysis (*A, C* and *E*) was evaluated via comparing our data against 1000 bootstrap tests, with * indicating p<0.05. One-way ANOVA was performed to evaluate statistical significance in gold labeling number data (*B, D* and *F*), with * indicating p<0.05. Statistical significance of EGFR population distribution between untreated and DCA-treated conditions (*G*-*I*) was evaluated using one-way ANOVA with * indicating p<0.05.

We found that DCA markedly enhanced PA nanoclustering and disrupted PS clustering, while having little effect on PIP_2_, PIP_3_ and cholesterol (Fig. 1A). By counting the number of the gold nanoparticles per 1μm^2^ PM area, we estimated changes in level of each type of lipids in the PM. DCA markedly elevated the level of PA but decreased the level of PIP_2_ in the PM of Caco-2 cells (Fig. 1B). Changes in the PM lipid levels were unlikely to be caused by alterations in lipid metabolism because of the short DCA treatment time; DCA incubation of at least 1h is required to induce any meaningful change in lipid metabolism [22]. Further, DCA caused no change in cholesterol aggregation pattern and PM level (Fig. 1A and B), consistent with the view that changes lipid levels observed in Fig. 1 were independent of lipid metabolism.

Conjugated bile acids are typically more hydrophilic than their unconjugated cognates, thus possessing less ability to influence PM lipids [1]. We then tested how taurine-conjugated DCA, taurodeoxycholic acid (TDCA), potentially influenced the spatial segregation of PA. We focused specifically on PA because DCA-induced elevation in PA clustering was the most striking. Fig. S1D shows that TDCA had minimal effect on the clustering of PA. Interestingly, TDCA treatment induced significant mis-localization of PA from the PM of Caoc-2 cells (Fig. S1E). Ursodeoxycholic acid (UDCA) and its conjugated cognate tauroursodeoxycholic acid (TUDCA) have been shown to be cytoprotective [1]. Fig. S1D and E show that either UDCA or TUDCA had no effect on PA clustering, but mis-localized PA from the PM. To further examine the well-documented cytoprotective effects of UDCA against DCA, we conducted similar EM-spatial analysis using Caco-2 cells treated with DCA alone, UDCA alone or combination of both DCA and UDCA. Fig. S1F and G show that co-treatment with 1μM DCA and 1μM UDCA completely abolished the enhancing effects of DCA alone on PA clustering and PM localization.

To further explore the dynamics of the DCA effect on PA, we conducted EM-spatial analysis in a time course experiment. DCA at 1μM enhanced PA nanoclustering and PM localization within 30 seconds, achieved a peak effect within 5 minutes, and returned to the baseline within 20 minutes (Fig. 1C and D). We next conducted dose response experiments by incubating Caco-2 cells with different concentrations of DCA (0.05 - 30μM) for 5 minutes. PA nanoclustering and PM level were elevated at 0.05μM and plateaued at ∼1μM DCA (EC_50_ ∼ 0.6μM, Fig. 1E and F).

### Bile acid enhances EGFR dimerization/oligomerization

PA is a major structural component of EGFR signaling nanoclusters and mediates EGFR-MAPK signal transmission [13, 15, 23]. DCA robustly activates EGFR-dependent MAPK signaling in many GI cell lines [22, 24, 25]. We then used EGFR as an example to examine how the DCA-induced changes in PM lipids potentially contribute to the function of the surface receptors. EM-spatial analysis revealed that DCA at 1μM enhanced EGFR-GFP nanoclustering and PM localization within 30 seconds, an effect that peaked at ∼5 minutes (Fig. 1C and D). DCA at low doses (EC_50_ ∼ 0.3μM and plateau at ∼1μM) robustly elevated EGFR nanoclustering and PM localization (Fig. 1E and F). Because EGFR dimerization/oligomerization is an essential step in EGFR signaling [7, 8], we further interrogated our spatial data and evaluated the potential effects of DCA on population distribution of EGFR in the PM. DCA at 1μM for 5 minutes decreased the EGFR monomer population, while significantly elevating the dimer and the oligomer populations of EGFR (Fig. 1G-I). Thus, effects of DCA on EGFR nanoclustering correlated well with its effects on PA clustering in both the time course and dose response studies.

### DCA enhances co-localization between EGFR and PA in Caco-2 basal PM

To further establish causality, we quantified the co-localization between the acidic lipids and EGFR in the Caco-2 basal PM using the bivariate EM co-localization analysis. The basal PM of Caco-2 cells co-expressing one of the GFP-tagged lipid binding domains and EGFR-RFP were attached to the EM grids and co-immunolabeled with both 6nm gold nanoparticles linked to anti-GFP antibody and 2nm gold nanoparticles coupled to anti-RFP antibody. The spatial distribution of the two populations of gold particles was imaged using TEM and analyzed using the bivariate K-functions (Fig. S2A and B). Fig. S2C shows the extent of co-localization, *L*_*biv*_(*r*)-*r*, plotted against the length scale, *r*. The *L*_*biv*_(*r*)-*r* values above the 95%C.I. of 1 indicate the statistically significant co-localization between the 6nm and 2nm gold populations. To summarize the co-localization data, we calculated the area-under-the-curve for the *L*_*biv*_(*r*)-*r* curves and termed as the L-bivariate integrated, or LBI. The high LBI values indicate more extensive co-localization while the LBI values <100 indicate no co-localization between the two gold populations [20, 21]. Fig. 2A shows the LBI values for the extent of co-localization between EGFR and various lipids. Interestingly, in the untreated Caco-2 cells, EGFR co-localized extensively with cholesterol (Fig. 2A); DCA at 1μM for 5 minutes markedly elevated the co-localization between EGFR and PA, while depleting cholesterol from the EGFR nanoclusters (Fig. 2A). DCA had little effect on the association of EGFR with other lipids tested, including PS, PIP_2_ and PIP_3_. Taken together, our data suggest that DCA has distinct local effects on the lipid composition of EGFR nanoclusters in the PM.

**Figure 2.**
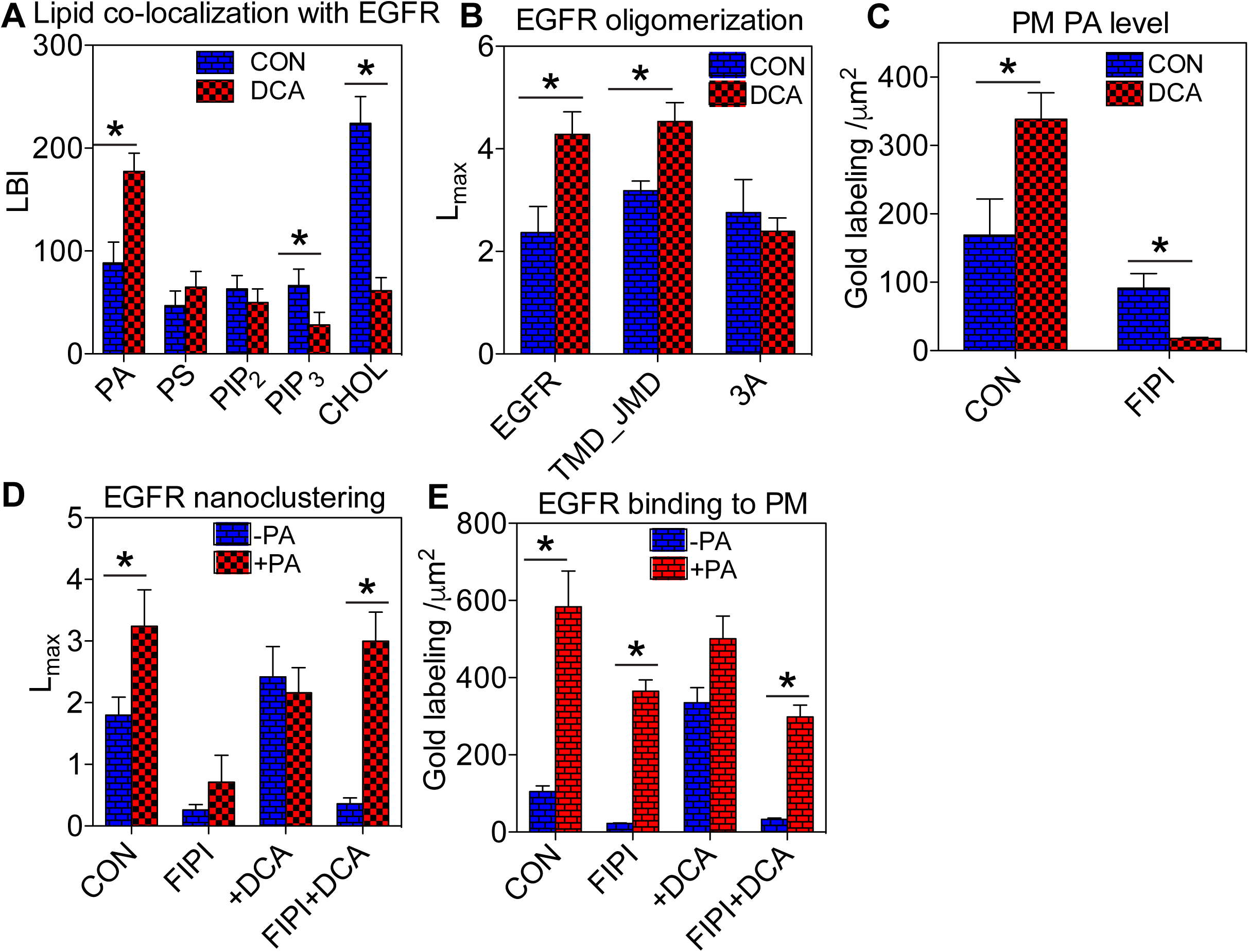
DCA-induced elevation of EGFR oligomerization in Caco-2 PM is mediated by PA. (**A**) Co-localization between various lipid-binding domains and EGFR was quantified using EM-bivariate co-localization analysis. Caco-2 cells co-expressing a GFP-tagged lipid-binding domain and EGFR-RFP were grown to a monolayer before treatment of 1μM DCA for 5 minutes. Intact basal PM of Caco-2 cells was attached to EM grids and immunolabeled with 6nm gold nanoparticles conjugated with anti-GFP antibody and 2nm gold linked to anti-RFP antibody, respectively. Gold distribution was imaged using TEM at 100,000x magnification and analyzed using bivariate K-function. LBI values indicate extent of co-localization between each lipid type and EGFR, with values >100 indicating statistically significant co-localization. (**B**) EM-univariate spatial analysis shows the extent of oligomerization of the full-length EGFR-GFP, the truncated transmembrane domain/juxtamembrane domain TMD_JMD-GFP, and the truncated mutant TMD_JMD_3A-GFP in the basal PM of Caco-2 cells without / with 1μM DCA for 5 minutes. (**C**) Caoc-2 cells expressing GFP-PASS (specifically binding to PA) were pre-treated with 0.75μM FIPI (pan-PLD inhibitor) for 25 minutes before co-incubation with both 0.75μM FIPI and 1μM DCA for an additional 5 minutes. In the same EM-spatial analysis, the number of gold particles with a 1μm^2^ PM area was counted to estimate changes in PA level in the cell PM. The same PA depletion experiments using FIPI were conducted on Caco-2 cells expressing EGFR-GFP without / with supplementation of 10μM exogenous egg PA. Oligomerization (**D**) and number of gold particles (**E**) within a 1μm^2^ PM area were quantified using EM-univariate clustering analysis. For all the EM-immunogold labeling univariate and bivariate experiments, at least 15 PM sheets from individual cells were analyzed in each condition. In all univariate and bivariate spatial analyses (*A, B, C*, and *E*), statistical significance between untreated controls and various treatments in clustering analyses was evaluated using bootstrap tests, with * indicating p<0.05. Statistical significance of gold numbers (*D* and *F*) was evaluated using one-way ANOVA, with * indicating p<0.05.

EGFR interacts with lipids in the PM via a single-span transmembrane domain (TMD) and an adjacent intracellular juxtamembrane domain (JMD) enriched with basic residues [7, 26]. To further examine whether the DCA-enhanced EGFR oligomerization is a membrane-mediated effect, we ectopically expressed a GFP-tagged truncated EGFR containing only the TMD and the JMD (amino acids 622-663), termed as TMD_JMD-GFP, in the Caco-2 cells. In the absence of all the extracellular domains and the intracellular phosphorylation sites, TMD_JMD-GFP is likely only sensitive to changes in the nano-environment of the PM. We then compared the ability of DCA to alter the oligomerization pattern of the full-length EGFR-GFP vs. the truncated TMD_JMD-GFP by performing the immunogold EM-univariate spatial analysis. Fig. 2C shows that, in the basal PM of the untreated Caco-2 cells, TMD_JMD-GFP clustered as efficiently as the full-length EGFR-GFP. The clustering of TMD_JMD-GFP also responded to DCA (1μM, 5 minutes) as effectively as its full-length cognate (Fig. 2B). Upon dimerization, the helical structure of the EGFR JMD of each monomer associates extensively with the anionic lipids in the PM using exposed basic residues, such as Arginine (Arg) 651, Arg657 and Arg662 [9]. The basic residues Arg645-647 also participate in binding with the anionic lipids in the membranes [6]. We, therefore, mutated Arg647, Arg651 and Arg662 to the neutral alanines to generate a triple alanine mutant TMD_JMD_3A-GFP. In an EM-univariate analysis, TMD_JMD_3A-GFP no longer responded to the DCA stimulation (Fig. 2B). Thus, our data suggests that EGFR JMD association with the anionic lipids in the PM mediates DCA-induced EGFR oligomerization.

To further investigate the role of PA in the DCA-enhanced EGFR nanoclustering, we pre-treated the Caco-2 cells with 5-fluoro-2-indolyl des-chlorohalopemide (FIPI), a pharmacological inhibitor of phospholipase D (PLD) that catalyzes the conversion of phosphatidylcholine (PC) to PA [15], before co-incubation with 1μM DCA. The FIPI treatment effectively abolished the ability of DCA to elevate PA levels in the Caco-2 PM (Fig. 2C). The FIPI treatment also disrupted EGFR oligomerization and PM localization and effectively abolished the DCA-induced elevation of EGFR oligomerization/PM localization (Fig. 2D and E). Supplementation with the exogenous egg PA effectively reversed the FIPI inhibitory effects (Fig. 2D and E). Our data suggest that the lipid messenger PA mediates the DCA-induced changes in EGFR oligomerization in the Caco-2 PM.

### PA mediates DCA-stimulation of EGFR-MAPK signaling

We then quantified the signaling effects of DCA on EGFR signaling. In a time-course study, the Caco-2 cells were serum-starved and then incubated with 1μM DCA for various time periods before harvesting. Western blotting was then used to quantify the ability of DCA to stimulate the phosphorylation of MEK/ERK in the MAPK pathway, as well as the phosphorylation of Akt in the PI3K pathway. Fig. 3A shows that DCA robustly elevated the levels of pMEK and pERK in the Caco-2 cells within 2-5 minutes, which decayed to the baseline within 15 minutes. This robust activation is consistent with the previous findings [22, 24, 25], and is also consistent with the time-course of DCA-elevated oligomerization of the EGFR (Fig. 1C and D). In a dose response study, the MAPK signal output was elevated by 50nM DCA and plateaued at ∼1μM with an EC50 of ∼0.2μM (Fig. 3B). The effective DCA doses for stimulating the EGFR-MAPK signaling were significantly lower than previously shown, but consistent with the effective DCA doses required to elevate the oligomerization of EGFR (Fig. 1E and F). DCA did not change the pAkt level in either the time course or the dose experiments, suggesting that DCA has no measurable effect on the PI3K pathway in Caco-2 cells.

**Figure 3.**
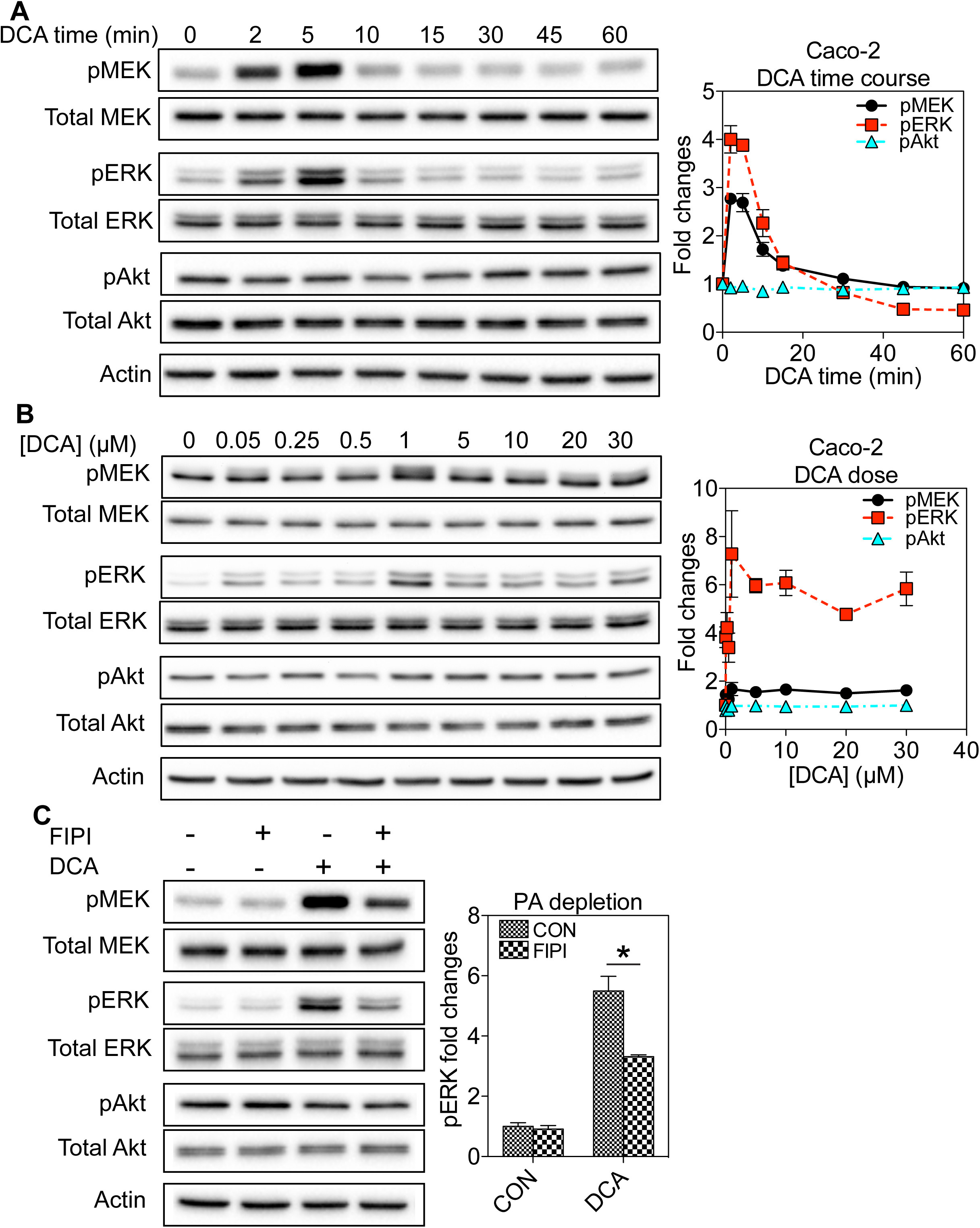
DCA stimulation of EGFR-MAPK signaling in Caco-2 cells is mediated by PA. (**A**) Caco-2 cells grown to a monolayer were pre-serum-starved before incubation with 1μM DCA for various time points to ensure that the total serum starvation time was 3 hours without / with DCA. Whole cell lysates were collected and blotted using antibodies against pMEK, total MEK, pERK, total ERK, pAkt or total Akt, as well as the loading control actin. (**B**) Caco-2 cells grown to a monolayer were pre-serum-starved for 2 hours and 55 minutes before incubation with various concentrations (1-30μM) DCA for 5 minutes. Whole cell lysates were collected and blotted using antibodies against pMEK, total MEK, pERK, total ERK, pAkt or total Akt, as well as the loading control actin. (**C**) Caco-2 cells were pre-serum-starved for 2 hours and 30 minutes before incubation with 0.75μM FIPI for 25 minutes and subsequent co-incubation with FIPI and 1μM DCA for 5 minutes. Whole cell lysates were collected for Western blotting against pMEK, total MEK, pERK, total ERK, pAkt or total Akt, as well as the loading control actin. In all signaling experiments, 3 individual experiments were performed separately and quantitation is shown as mean±SEM. Statistical significance was evaluated using one-way ANOVA, with * indicating p<0.05.

The essential role of EGFR in the DCA-induced stimulation of the MAPK signaling has been shown before [22, 24, 25]. We also validated this via pre-treating the Caco-2 cells with a specific EGFR inhibitor AG1478 before the DCA exposure. The AG1478 pre-treatment completely abolished the stimulatory effects of DCA (Fig. S2D), showing that DCA specifically targets EGFR. Concordantly DCA had no effect on the MAPK signaling in the wild-type Chinese hamster ovarian (CHO) cells, which do not possess the endogenous EGFR [27], but effectively elevated the MAPK signaling in the CHO cells stably expressing EGFR-GFP (Fig. S2E). These data clearly show the essential role of EGFR in the DCA-stimulation of the mitogenic signaling. Depletion of the endogenous PA via the FIPI treatment significantly abrogated the ability of DCA to stimulate the EGFR/MAPK signaling (Fig. 3C). Taken together, these data suggest that generation of PA mediates the DCA stimulation of the EGFR-MAPK pathway.

### PA mediates DCA stimulation of EGFR signaling in ex vivo human intestinal enteroids

To further examine the physiological relevance of PA in mediating the effects of DCA, we conducted signaling experiments using *ex vivo* human intestinal enteroids. We first conducted the time-course and the dose response experiments using the human intestinal enteroids; similar to Caco-2 cells, DCA rapidly elevated the MAPK signaling which peaked at 5 minutes and returned back to the baseline within 15 minutes (Fig. 4A). The DCA dose response plateaued at ∼5μM (Fig. 4B). Pre-treatment with the pan-PLD inhibitor FIPI effectively abolished the ability of DCA to activate the MAPK signaling (Fig. 4C). Taken together, DCA had a similar ability to enhance the PA-mediated EGFR-MAPK signaling in the *ex vivo* human tissue.

**Figure 4.**
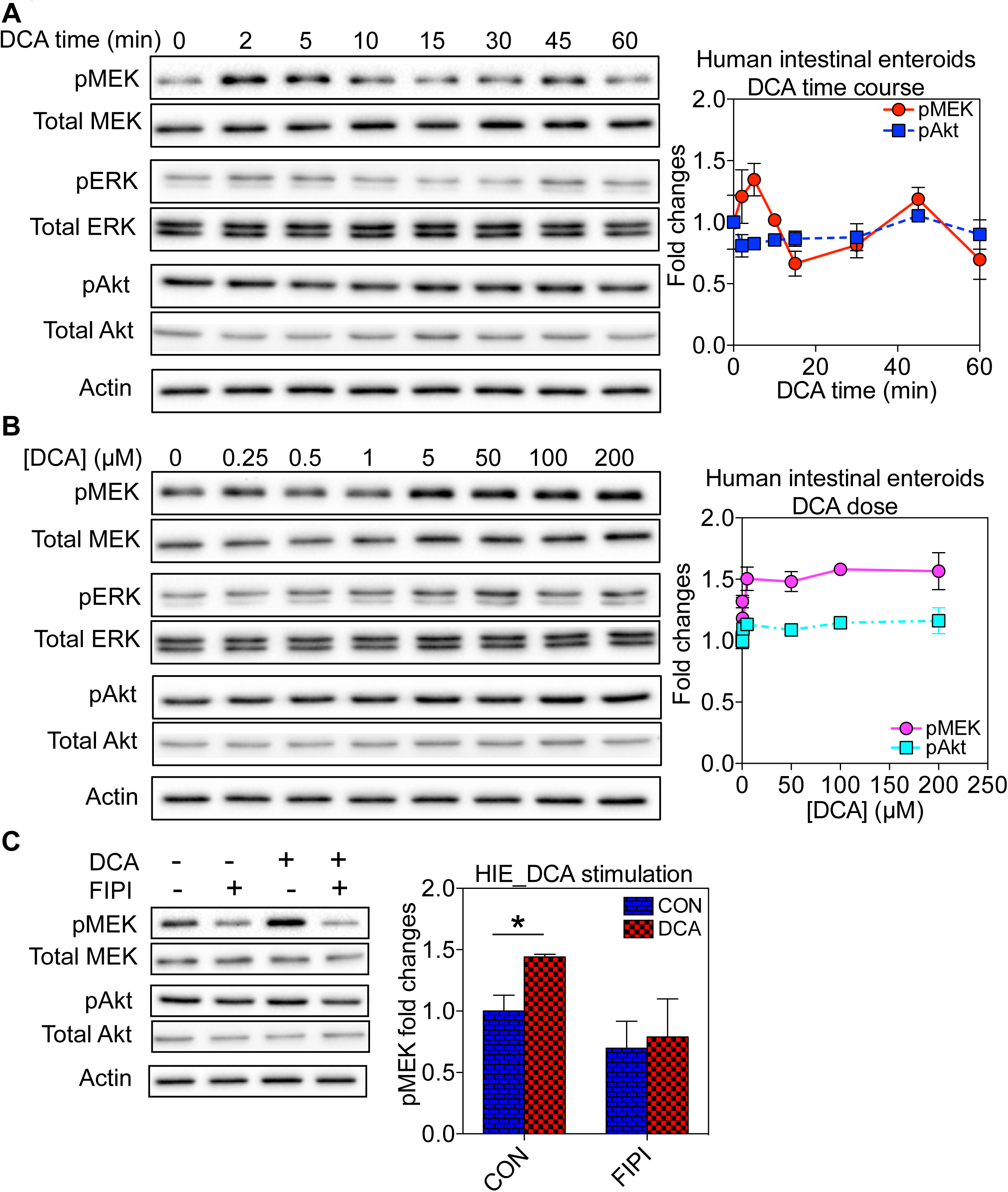
DCA stimulation of EGFR-MAPK signaling in *ex vivo* human intestinal enteroids is mediated by PA. (**A**) Human intestinal enteroids were pre-serum-starved before incubation with 5 μM DCA for various time points to ensure total serum starvation time is 2 hours for all conditions. Whole cell lysates were collected and blotted using antibodies against pMEK, total MEK, pERK, total ERK, pAkt or total Akt, as well as the loading control actin. (**B**) Human intestinal enteroids were serum-starved for 1 hour and 55 minutes before incubation with 1-200μM DCA for an additional 5 minutes. Whole cell lysates were collected and blotted using antibodies against pMEK, total MEK, pERK, total ERK, pAkt or total Akt, as well as the loading control actin. (**C**) Pre-serum-starved human intestinal enteroids were treated with 0.75μM FIPI for 25 minutes and subsequent co-incubation with both FIPI and 5μM DCA. Whole cell lysates were collected and blotted using antibodies against pMEK, total MEK, pERK, total ERK, pAkt or total Akt, as well as the loading control actin. In all signaling experiments, 3 individual experiments were performed separately and quantitation is shown as mean±SEM. Statistical significance was evaluated using one-way ANOVA, with * indicating p<0.05.

### DCA modulates lipid dynamics and cell signaling in non-GI cells in a similar manner

Low levels of the bile acids (<5μM), of which ∼30% can be DCA, can be found in the circulation [28]. Our data showing that DCA at doses <1μM robustly enhanced EGFR oligomerization and signaling in the GI cells and tissues suggest that a similar behavior can occur in non-GI cells/tissues. To test this, we first conducted the time-course and the dose studies to examine how DCA potentially influence the EGFR-MAPK and the PI3K cascades in the baby hamster kidney (BHK) cells. The time-course for the DCA stimulation of the MAPK pathway in the BHK cells was similar to the Caco-2 cells (Fig. S3A). Strikingly, DCA effectively decreased the pAkt levels by >50% within 5 minutes (Fig. S3A), suggesting effective inhibition of the PI3K pathway by DCA in the BHK cells. We also validated the EGFR-dependence by pre-treating the BHK cells with AG1478. As expected, inhibiting the EGFR activity effectively abolished the DCA stimulation of the EGFR-MAPK signaling (Fig. 3C). To examine if DCA affected the EGFR oligomerization in the BHK cells, we conducted the EM-univariate clustering analysis. The spatial analysis showed that DCA enhanced the oligomerization of EGFR-GFP in the apical PM of the BHK cells (Fig. S3D). Taken together, our data suggest that DCA has the similar stimulatory effects on the EGFR/MAPK signaling in non-GI cells. The PI3K pathway, on the other hand, responded differently in the Caoc-2 cells vs. the BHK cells, suggesting the potential tissue-specific responses.

## Discussion

Our current study aims to explore potential molecular mechanism(s) for the bile acid-induced activation of the cell surface EGF receptor. We found that the secondary bile acid DCA has selective effects on different acidic lipids in the PM of the Caco-2 cells. Especially, DCA markedly enhances the local spatial aggregation of PA, which in turn induces the co-localization between PA and EGFR, promotes the EGFR dimerization/oligomerization and stimulates the EGFR-MAPK signaling. Acute PA depletion effectively abolishes the DCA-induced EGFR oligomerization and the DCA stimulation of the EGFR-MAPK signaling. This effect has been observed in both the cultured human colon cancer cell line and the *ex vivo* human intestinal enteroids. Thus, the dynamic nano-domains of the anionic lipids in the PM may act as the signaling platforms to mediate the bile acid stimulation of the cell surface receptors.

Dependence of the EGFR oligomerization on the acidic lipids has been demonstrated before. The EGFR likely engages with PM lipids via the hydrophobic single-span TMD and the adjacent intracellular JMD enriched with positively charged residues [6, 13, 29]. Indeed, a truncated EGFR fragment containing only the TMD and JMD clusters as efficiently as the full-length EGFR in the PM (Fig. 2C). Binding to the acidic lipids via the polybasic sequences in its JMD drives allosteric folding of the EGFR cytoplasmic domain, which is a key step in the EGFR dimerization/oligomerization and the consequential auto-phosphorylation and the activation [6, 13, 29]. Specifically, EGFR preferentially associates with PA and depends on PA interactions for the oligomerization and the signal transduction [13]. Concordantly, mutating the basic residues in the JMD significantly comprises the ability of DCA to elevate EGFR oligomerization and PM binding (Fig. 2G). Thus, altering the local spatial distribution of PM lipids surrounding EGFR potentially changes EGFR activities. By contrast, PS, also a monovalent anionic phospholipid, has no effect on the EGFR oligomerization [13], suggesting that EGFR possesses specific lipid sorting capacity rather than simply sensing global electronegativity on the PM.

It is also interesting that DCA induces local cholesterol depletion from EGFR, while EGFR co-localizes with cholesterol extensively in the untreated Caco-2 cells. The role of cholesterol in regulating the EGFR activation is controversial. While some studies show that cholesterol inhibits the EGFR ligand-independent activation [26], others propose that cholesterol positively regulates the EGFR stimulation [30]. Our current data suggest that cholesterol content surrounding EGFR is highly transient. While EGFR co-localizes with cholesterol extensively at the basal state, cholesterol is depleted from EGFR upon the DCA incubation. Although our data is consistent with the inhibitory effects of cholesterol on EGFR, we cannot rule out the possibility that cholesterol enrichment at the basal state is still required for its proper activation.

It is not entirely clear how DCA specifically targets PA in the PM. Amphipathic bile acids, such as DCA, possess the capability to intercalate in the PM and shift the lateral distribution of the lipids. Indeed, recent biophysical studies show that DCA at low concentrations well below its CMC preferentially partitions into highly fluid liquid-disordered domains and alters the lateral spatial segregation of the phospholipids in synthetic and isolated native membranes [10, 11]. DCA <15μM also induces marked changes in the cross-sectional area of lipids in monolayers [12]. These *in vitro* studies show that DCA is capable of changing the dynamic local distribution of the lipids in the membranes. Thus, the lateral distribution of PA lipids may be altered as a result of DCA-induced shift in heterogeneity of the PM. Furthermore, the DCA treatment may also alter the lipid recycling, which leads to the observed changes in overall PA levels in the PM in our EM immunogold labeling analysis. This is indeed supported by our EM data on the PM localization of EGFR. The stimulation of EGFR results in the internalization of the receptor, which coincides with the changes in the level of PA in the PM.

In addition to PA, DCA also significantly disrupts PS clustering and mislocalizes PIP_2_ from the basal PM of the Caco-2 cells. Although not clear of the underlying mechanisms, these effects may influence cell functions dependent on these lipids. For instance, PIP_2_ regulates or inhibits functions of many ion channels on the cell PM. Particularly, many cholestatic patients with elevated serum bile acid levels suffer debilitating itch [31], mechanism of which is still unknown. Transient receptor potential Ankyrin sub-family 1 (TRPA1) is the ion channel mostly implicated in chronic itching [32]. PIP_2_ associates with TRPA1 and inhibits its channel function [33]. Thus, DCA can potentially activate TRPA1 channels and cause chronic itching partially via depleting local PIP_2_ level, although bile acid specific receptor TGR5 has also been implicated [34].

## Conclusion

Here we propose that the secondary bile acid, DCA, stimulates EGFR-MAPK signaling via modulating the dynamic spatial distribution of minor acidic lipids in the PM. DCA has distinct effects on different PM lipids, which potentially leads to highly diverse responses from various lipid-dependent surface receptors, ion channels and other membrane-associating proteins. This lipid-mediated effect potentially contributes to the diverse biological and pathophysiological effects of bile acids.

## Experimental procedures

### Electron microscopy (EM)-spatial analysis

#### EM-univariate clustering analysis

The univariate spatial analysis quantifies the extent of nanoclustering of a single species of immunogold-labeled constituent on cell PM. Caco-2 cells ectopically expressing GFP-tagged lipid-binding domains or EGFR were seeded on gold EM grids and grown to a monolayer. Apical portion of the cells was peeled off using a filter paper wetted with PBS, leaving only intact basolateral PM attached to the gold EM grids. Basal PM was then fixed in 4% paraformaldehyde (PFA) and 0.1% gluaraldehyde, immunolabeled with 4.5nm gold nanoparticles conjugated to anti-GFP antibody and negative-stained with uranyl acetate. Gold particles on intact basal PM was imaged using TEM at 100,000x magnification. The coordinates of each gold particle were assigned using ImageJ. Spatial distribution of gold particles was quantified using Ripley’s K-function, which tests the null hypothesis that all points in a selected area are distributed randomly (Eqs. 1 and 2):

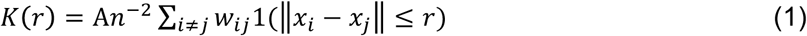

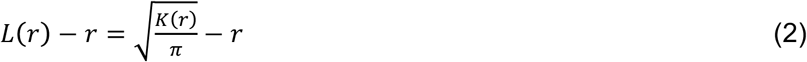

where *K(r*) = univariate K-function for a distribution of *n* gold particles in a basal PM area of *A*; *r*= length scale between 1 and 240 nm at 1nm increments ‖·‖ = Euclidean distance; indicator function of 1(·) is assigned a value of 1 if ‖ *x*_*i*_-*x*_*j*_‖ ≤ r and is assigned a value of 0 if ‖*x*_*i*_-*x*_*j*_‖ < r. An unbiased edge correction for points at the edge of the study area was achieved via including *W*_*ij*_^-1^ = the proportion of the circumference of a circle that has the center at *x*_*i*_ and radius ‖ *x*_*i*_-*x*_*j*_‖. *L*(*r*) - *r* is a linear transformation of *K*(*r*) and is normalized against the 99% confidence interval (99% C.I.) estimated from Monte Carlo simulations. Points distributed in a complete random manner yields a *L*(*r*) - *r* value of 0 for all values of *r*. On the other hand, spatial clustering at certain length scale yields a *L*(*r*) - *r* value above the 99% C.I. of 1 at the corresponding value of *r*. At least 15 PM sheets were imaged, analyzed and pooled for each condition in the current study. Statistical significance was evaluated via comparing our calculated point patterns against 1000 bootstrap samples in bootstrap tests [35].

#### EM-Bivariate co-localization analysis

The bivariate co-localization analysis quantifies co-clustering / co-localization between GFP-tagged and RFP-tagged of proteins / peptides on the PM inner leaflet. This bivariate analysis allows quantification of co-localization over a wide range of distances between 8-240nm. Intact basal PM of Caco-2 cells co-expressing both populations of proteins was attached and fixed to EM grids. GFP- and RFP-tagged proteins on the intact basal PM sheets were immunolabeled with 2nm gold conjugated to anti-RFP antibody and 6nm gold linked to anti-GFP antibody. X and Y coordinates to each population of gold particles were assigned using ImageJ. Gold particle co-localization was calculated using a bivariate K-function, which tests the null hypothesis that the two point populations spatially segregate from each other. (Eqs. 3-6):

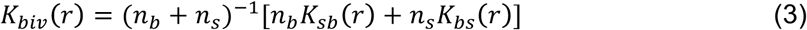

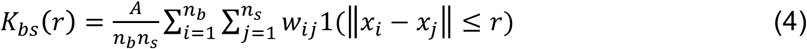

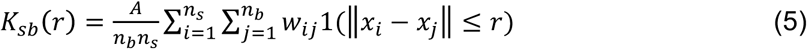

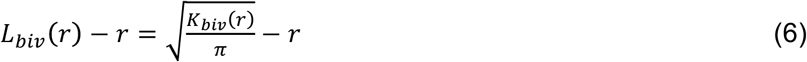

where *K*_*biv*_(*r*) = the bivariate estimator composed of 2 individual bivariate K-functions: *K*_*bs*_(*r*) for the distribution of the 6nm-big gold particles (*b* = big gold) with respect to each 2nm-small gold particle (*s* = small gold); and *K*_*sb*_(*r*) for the distribution of the small gold particles with respect to each big gold particle. An intact PM sheet with an area *A* contains n_b_, number of 6nm big gold particles and n_s_, number of 2nm small gold particles. Other notations are the same as in Eqs. A and B above. *L*_*biv*_(*r*)-*r* is a linear transformation of *K*_*biv*_(*r*), and was normalized against the 95% confidence interval (95% C.I.) estimated from Monte Carlo simulations. Spatial segregation between the two populations of gold particles yields an *L*_*biv*_(*r*)-*r* value of 0 for all values of r. On the other hand, co-localization that occurs at certain distance yields an *L*_*biv*_(*r*)-*r* value above the 95% C.I. of 1 at the corresponding distance of *r*. Each *L*_*biv*_(*r*)-*r* curve was then integrated to yield a value of area-under-the-curve over a fixed range 10 < *r* < 110 nm, which was termed bivariate *L*_*biv*_(*r*)-*r* integrated (or LBI):

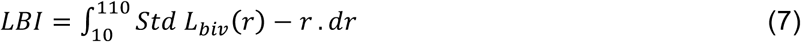

For each condition, at least 15 basal PM sheets were imaged, analyzed and pooled to show as mean of LBI values ± SEM. Statistical significance between conditions was evaluated via comparing against 1000 bootstrap samples as described [35].

#### Cultured cells and Western blotting

Caco-2 cells were grown in EMEM containing 20% FBS until a monolayer formed. BHK cells were grown in DMEM containing 10% BCS until 85-90% confluency. CHO cells were grown DMEM containing 10% FBS until 85-90% confluency. All cultured cells were pre-serum-starved before DCA treatment in all experiments. In all time course, dose dependence and inhibition studies, total serum starvation time was kept the same for all conditions for each cell line. Specifically, total serum starvation time was 3 hours for Caco-2 cells, 2 hours for BHK cells and 2 hours for CHO cells.

Whole cell lysates were collected and blotted using antibodies against pMEK, pERK and pAkt. Actin was used as loading control. Each experiment was performed at least 3 times separately and data was shown as mean±SEM. Statistical significance was evaluated using one-way ANOVA, with * indicating p < 0.05.

#### Human intestinal enteroids

Handling of human intestinal enteroids was the same as described before [36]. Briefly, de-identified human intestinal stem cells isolated from patients were collected and mixed with pre-chilled Matri gel, plated in 24-well plates and grown in complete medium with growth factors (CMGF+). Enteroids were allowed to mature for a week before experiment. On the day of experiments, enteroids were washed 3x in complete medium without growth factors (CMGF-) and incubated in CMGF-medium for partial serum starvation. Compositions of CMGF+ and CMGF+ media followed exactly those in [36]. For all conditions, the total serum starvation time was 2 hours including each type of treatment.

Whole cell lysates were collected and blotted using antibodies against pMEK, pERK and pAkt. Actin was used as loading control. Each experiment was performed at least 3 times separately and data was shown as mean±SEM. Statistical significance was evaluated using one-way ANOVA, with * indicating p < 0.05.

## Acknowledgments

This work is partially supported by and the National Institutes of Health (NIH: P30 DK056338) and the Cancer Research and Prevention Institute of Texas (CPRIT) (RP130059 and RP170233).

## Supporting Information

**Supplemental Figure 1. EM univariate clustering analysis quantifies extent of oligomerization of lipids and EGFR in Caco-2 basolateral PM.** EM micrographs of intact basolateral PM of Caco-2 cells ectopically expressing GFP-PASS under control condition (**A**) or treated with 1μM DCA for 5 minutes (**B**). GFP-PASS bound to PA on the intact Caco-2 basal PM was immunolabeled with 4.5nm gold nanoparticles conjugated with anti-GFP antibody. (**C**) Univariate clustering of gold particles labeling various GFP-tagged lipid-binding domains without / with DCA (1μM, 5 minutes) was calculated using K-function analysis. Extent of clustering, *L*(*r*)-*r*, was plotted against length scale, *r*, with the peak value termed as *L*_*max*_. *L*(*r*)-*r* values above the 99% confidence interval (99%C.I.) of 1 indicate statistically meaningful clustering, whereas *L*(*r*)-*r* values below 99%C.I. indicate uniform distribution. Optimal clustering, *L*_*max*_, (**D**) and gold labeling (**E**) of GFP-PASS in Caco-2 cells treated with 1μM of various unconjugated and tauro-conjugated bile acids were quantified using EM-univariate clustering analysis. Optimal clustering (**F**) and PM binding (**G**) of GFP-PASS in Caoc-2 cells exposed to DCA alone, UDCA alone or combination of both DCA and UDCA were quantified using EM-univariate spatial analysis. Statistical significance between untreated and DCA-treated conditions in all clustering analyses was evaluated using bootstrap tests, with * indicating p<0.05. Statistical significance between untreated and DCA-treated conditions in gold labeling was examined using one-way ANOVA, with * indicating p<0.05.

**Supplemental Figure 2. EM-bivariate co-localization analysis quantifies extent of lipid enrichment surrounding EGFR in Caoc-2 basal PM.** EM micrographs of intact basal PM of Caco-2 cells co-expressing GFP-PASS and EGFR-RFP untreated (**A**) or treated with 1μM DCA for 5 minutes (**B**). GFP-PASS was immunolabeled with 6nm gold conjugated to anti-GFP antibody, while EGFR-RFP was immunolabeled with 2nm gold coupled to anti-RFP antibody. (**C)** Bivariate K-function calculated the co-localization between 6nm and 2nm gold populations on intact Caco-2 basal PM sheets. Extent of co-localization, *L*_*biv*_(*r*)-*r*, was plotted against length scale, *r*. *L*_*biv*_(*r*)-*r* values above the 95%C.I. indicate statistically significant co-localization. Each *L*_*biv*_(*r*)-*r* curve was then integrated between *r* values of 10 and 110 to yield integrated L-bivariate, or LBI, to summarize the spatial data. Statistical significance between untreated and DCA-treated conditions in bivariate co-localization analyses was evaluated using bootstrap tests, with * indicating p<0.05. (**D**) Caco-2 cells grown to a monolayer were serum-starved for 2 hours and 30 minutes before treatment of 1μM EGFR specific inhibitor AG1478 for 25 minutes and a subsequent co-incubation with AG1478 and 1μC;M DCA for 5 minutes. Whole cell lysates were collected and blotted using antibodies against pMEK, pERK, or pAkt. (**E**) Wild-type CHO cells or CHO cells stably expressing EGFR-GFP were serum-starved for 10 minutes before incubation with 30μM DCA for various time points to ensure total serum starvation time is 30 minutes. Whole cell lysates were collected and blotted against pMEK. Statistical significance was evaluated using one-way ANOVA, with * indicating p<0.05.

**Supplemental Figure 3. DCA stimulates EGFR-MAPK signaling in non-GI cells in similar manner as Caoc-2 cells.** (**A**) BHK cells grown to 80-90% confluency were serum-starved for 1 hour before incubation with 30μM DCA for various time points to achieve total serum starvation time of 2 hours. Whole cell lysates were used to blot against pMEK, total MEK, pERK, total ERK, pAkt or total Akt, as well as the loading control actin. (**B**) BHK cells grown to 80-90% confluency were serum-starved for 1 hour and 55 minutes before incubation with various concentrations of DCA for 5 minutes. Whole cell lysates were used to blot against pMEK, total MEK, pERK, total ERK, pAkt or total Akt, as well as the loading control actin. (**C**) BHK cells were serum-starved for 1 hour and 30 minutes before treatment with 1μM AG1478 for 25 minutes and a subsequent co-incubation with AG1478 and 30μM DCA for 5 minutes. Whole cell lysates were used to blot against pMEK, pERK, or pAkt. Statistical significance was evaluated using one-way ANOVA, with * indicating p<0.05. (**D**) EM-univariate clustering experiment was conducted in BHK cells expressing EGFR-GFP without / with 30μM DCA. Intact apical PM sheets of non-polarized BHK cells were attached to EM grids and immunolabeled with 4.5nm gold particles conjugated to anti-GFP antibody. Univariate clustering of the gold particles was quantified using univariate K-function analysis. *L*_*max*_ values indicate the extent of oligomerization of EGFR-GFP in apical PM of BHK cells. Statistical significance between untreated and DCA-treated conditions in univariate clustering analyses was evaluated using bootstrap tests, with * indicating p<0.05.

